# Deep generative models generate mRNA sequences with enhanced translation capacity and stability

**DOI:** 10.1101/2024.06.20.599727

**Authors:** He Zhang, Hailong Liu, Yushan Xu, Yiming Liu, Jia Wang, Yan Qin, Haiyan Wang, Lili Ma, Zhiyuan Xun, Timothy K. Lu, Jicong Cao

## Abstract

Despite the tremendous success of messenger RNA (mRNA) COVID-19 vaccines, the extension of this modality to a broader spectrum of diseases necessitates substantial enhancements, particularly in the design of mRNAs with elevated expression levels and extended durability. Here we present GEMORNA, a deep generative model designed to generate novel mRNA coding sequences (CDSs) and untranslated regions (UTRs) with superior translation capacity, comparable to the sophisticated task of language translation and free-form poetry composition with accurate grammar and semantics. Our AI model was trained on an extensive collection of RNA sequences from diverse families, further enhanced with labeled data to refine its performance. Remarkably, we demonstrate that our AI-generated mRNAs exhibited 8.2-fold and 15.9-fold increases in firefly luciferase expression compared to benchmark mRNAs in two different cell types. Additionally, Our AI-designed COVID-19 mRNA vaccine elicited a 4-fold increase in anti-COVID antibody titer in mice relative to BNT162b2. Furthermore, GEMORNA’s versatility extends to circular mRNA design, which we facilitated a 27-fold increase in human erythropoietin protein expression *in vivo* than a systematically optimized benchmark sequence. We also created circular mRNAs with substantial improvements in expression levels, durability and anti-tumor cell cytotoxicity in mRNA-transduced CAR-T cells compared with an experimentally validated benchmark. In summary, GEMORNA generates novel mRNA sequences with significant performance improvements and has the potential to enable a wide range of therapeutic and vaccine applications.

## Introduction

Messenger RNA (mRNA) vaccines have established their potential as an effective approach to prevent severe COVID-19 disease. There are numerous efforts to extend mRNA therapeutics to other indications, including cancers and rare diseases (1; 2), but stronger and longer-lasting protein expression is necessary for these applications to be successful. This is challenging because mRNA designs exhibit wide differences in expression-related properties. For example, in mammalian cells, mRNA expression levels vary by 10^6^-fold, and half-lives vary by 40-fold (3) (**Fig. 1A**). mRNA sequence components, such as coding sequences (CDSs) and untranslated regions (UTRs), are crucial determinants of therapeutic activity (4). Thus, designing optimal mRNA sequences is of paramount importance to realizing the broad potential of mRNA therapeutics, but remains extremely challenging due to the extensive potential mRNA sequence space (**Fig. 1B**).

**Fig. 1:**
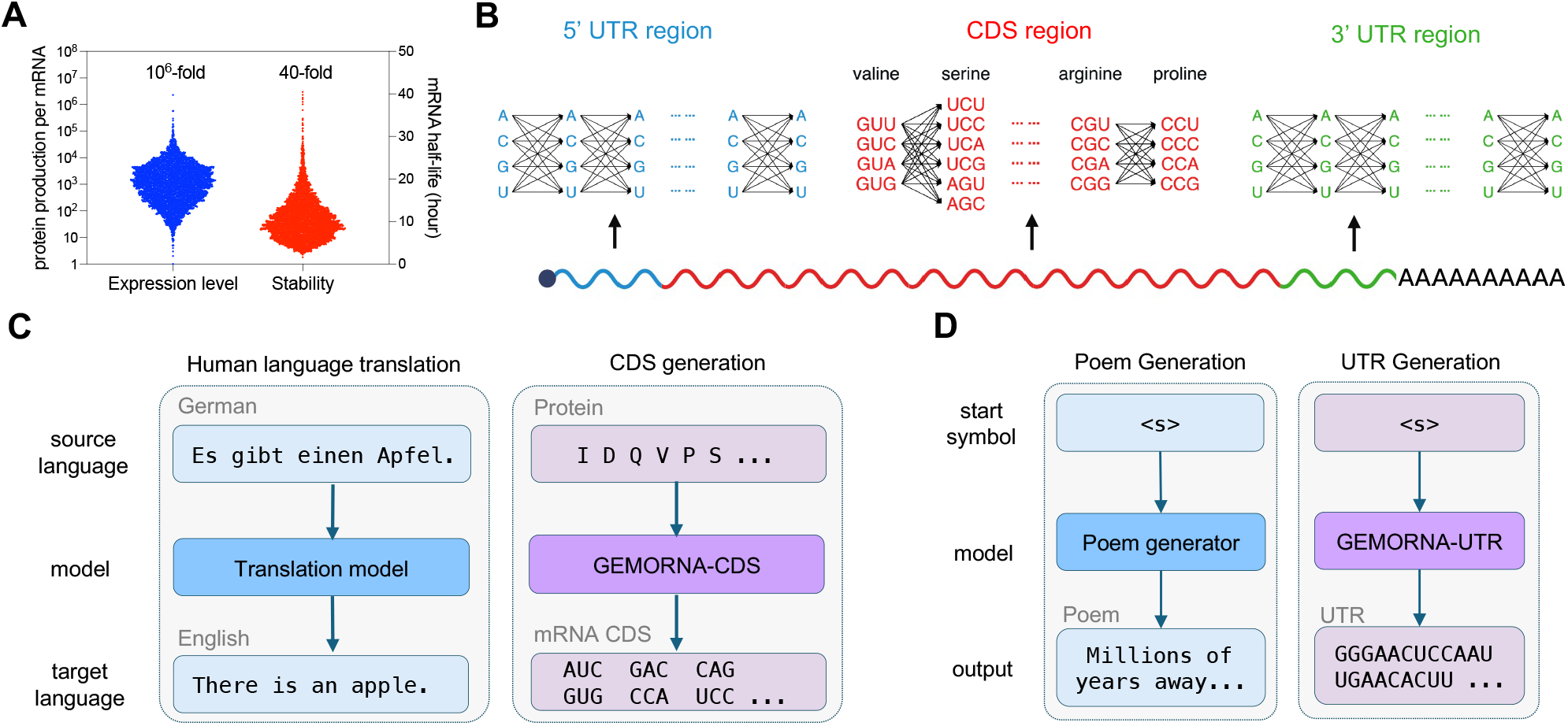
Overview of designing therapeutic mRNAs with GEMORNA. **A**) There are tremendous differences in gene expression levels and in-cell stability among 4,272 endogenous human mRNAs (3; 12). **B**) Schematic illustration of mRNA components, including the 5’ UTR, the CDS, and the 3’ UTR, resulting in a vast potential mRNA design space. Arrows between adjacent nucleotides (or codons) represent the possible choices at each position, and each path in the graph represents a different UTR (or CDS) sequence. **C**) Analogous to human language translation, GEMORNA-CDS transforms a protein sequence into a CDS sequence. **D)** Analogous to poem generation, GEMORNA-UTR generates UTR sequences from scratch.

In past decades, numerous efforts have sought to address the challenges of mRNA sequence optimization. Previous studies have demonstrated that optimizing codon usage within the CDS can enhance mRNA translation efficiency (5). A common approach involves optimizing the CDS codon adaptation index (CAI) by replacing rare codons with optimal synonymous codons (6). A more advanced strategy further incorporates codon pair usage (7). However, these methods remain tethered to local optimization of isolated codons or adjacent codons. Recently, deep learning-based models have been proposed for CDS optimization (8; 9). These models utilize Long Short-Term Memory (LSTM) networks, which lack an attention mechanism to extract long-dependencies in CDS sequences. Moreover, LSTM exhibits low efficiency due to non-parallelized training, which limits the size of the training data and hinders model generalizability. A recent study incorporated structural features, such as minimum free energy (MFE), into the optimization objective, enabling global optimization via dynamic programming (10). This algorithm led to improved stability and immunogenicity of chemically unmodified mRNAs, demonstrating a 128-fold increase in antibody response to COVID-19 mRNA vaccines compared to a codon-optimized baseline. However, this state-of-the-art algorithm has not been adapted for mRNAs containing modified nucleosides due to the lack of a free energy model inclusive of N1-methylpseudouridine (m1Ψ) (10), resulting in mismatch and reduced effectiveness for designing sequences of chemically modified mRNAs (4, 11).

It has been challenging to design 5’ UTRs de novo since the mechanisms governing 5’UTR regulation of mRNA translation is not fully understood. Thus, 5’ UTRs of natural mRNAs coding for highly abundant proteins, such as human alpha globin and human cytochrome B-245 alpha polypeptide (CYBA), have been used to drive synthetic gene expression. Recently, Zeng *et al*. developed a 5’ UTR design approach based on minimizing 5’ UTR secondary structure that demonstrated enhanced translational efficiency in cell-based assays (12). However, other factors that may influence initiation efficiency, such as mean ribosome load (MRL), were not considered. Several groups have described machine learning-based methods for 5’ UTR design (13; 14), which initially trained predictive models on labeled data and then incorporated them into genetic algorithms for 5’ UTR sequence evolution. These methods depend heavily on the reliability of the predictive models, and the evolutionary algorithms can be hindered by local optima. Similar to CDS optimization, current UTR design strategies only explore a minuscule fraction of the vast 5’ UTR design space, and most previous studies only compared to natural 5’ UTRs, while comprehensive comparisons with a diverse range of counterparts are still lacking. As a result, it remains unclear whether 5’ UTRs designed with these algorithms outperform the most advanced 5’ UTRs, such as those used in commercial products, and whether these algorithms can routinely discover high-performance and novel 5’ UTRs in the vast potential design space.

To overcome the limitations of previous methods, we created generative models to design mRNA sequences with improved translation capacity. Generative models have been used successfully for text generation. Drawing parallels between human language and genetic language, we adapted the concept and architecture of these generative models to mRNA sequences. Specifically, designing a CDS for a given protein is analogous to generating a translated sentence from its source language (**Fig. 1C**), and designing a UTR is akin to freely composing a poem due to the absence of protein sequence constraints (**Fig. 1D**). We developed our generative model GEMORNA, which stands for Generative models for RNA, as the core of our mRNA design process (**Fig. 2A**). We performed cell-based experiments to confirm that both GEMORNA-generated CDSs and UTRs achieve higher gene expression levels and durability compared to a diverse set of widely used controls, including natural, algorithm-optimized, and commercial sequences (**Fig. 2B**). By integrating multiple AI-generated RNA elements, we designed full-length therapeutic mRNAs with improved translation capability and immunogenicity compared with benchmark mRNAs and a marketed mRNA vaccine. Moreover, we demonstrated that in addition to linear mRNAs, GEMORNA can generate novel circular RNA sequences (circRNAs) with exceptional gene translation capacity. Finally, we used GEMORNA to create CD19 chimeric antigen receptor (CAR)-expressing circRNAs and showed that non-virally transduced T cells exhibited significantly stronger tumor-killing efficacy compared to a patented CD19-CAR circRNA benchmark. In summary, our generative AI models enable the de novo generation of full-length mRNAs with high gene expression levels and durability and have the potential to improve the potency of mRNA drugs across a variety of therapeutic areas.

**Fig. 2:**
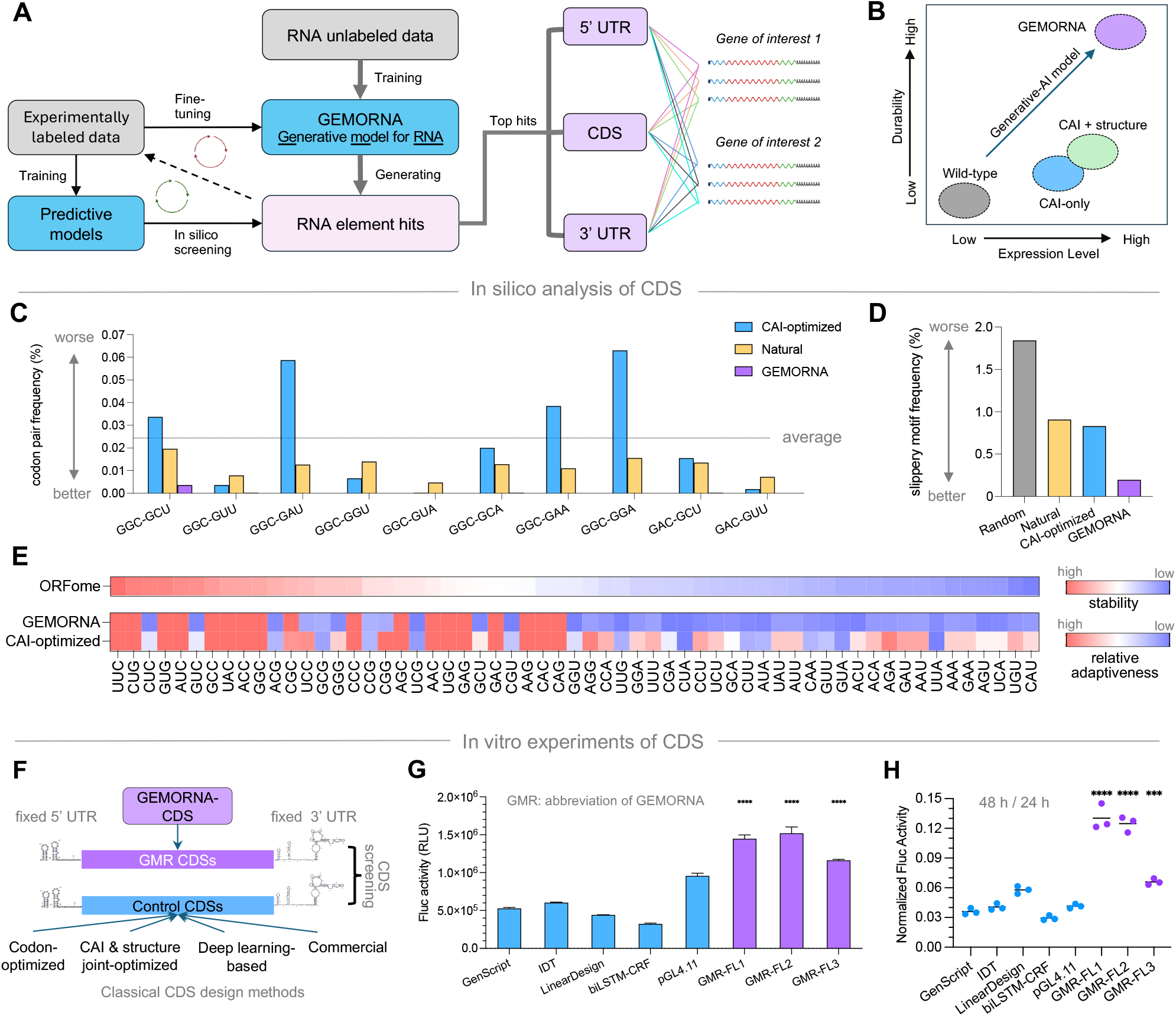
The architecture of our GEMORNA-CDS model for CDS design, along with *in silico* analysis and *in vitro* experimental results of CDS sequences generated by GEMORNA-CDS. Compared to conventionally designed CDS sequences, GEMORNA- generated CDSs exhibited better codon usage and enhanced translation capacity. **A)** GEMORNA-enabled mRNA design process. **B)** GEMORNA can de novo design therapeutic mRNAs with higher expression levels and durability versus previous rational design methods, such as those that use CAI (codon adaptation index), with or without structural information. **C)** Frequency of unwanted codon pairs in CAI-optimized, natural, and GEMORNA sequences; the dashed line is the mean frequency of all codon pairs. **D)** The frequency of slippery motifs in randomly generated, CAI-optimized, natural, and GEMORNA sequences. **E)** Codon stability coefficients measured by ORFome-assay (first row), and codon relative adaptiveness calculated from the codon frequencies of GEMORNA CDSs (second row) or CAI-optimized CDSs (third row). The relative adaptiveness of a codon is determined by dividing its frequency by the frequency of its most frequently occurring synonymous codon. **F)** Overview of the *in vitro* experiments for CDS validation. GEMORNA-generated CDSs (abbreviated as “GMR-”) were compared to a variety of CDSs generated from classical methods, while UTRs were kept unchanged. **G)** Expression levels of different Fluc2P CDSs at 24 h in HEK293T cells. **H)** Ratio of expression levels at 48 h to expression levels at 24 h in HEK293T cells line as a measure of in-cell stability over time. \*\*\**P <* 0.001, \*\*\*\**P <* 0.0001.

## Results

### GEMORNA-CDS: deep generative model for CDS element design

We first sought to develop generative models for CDS design, as CDSs are the core of mRNA therapeutics and vaccines. Therefore, we developed a GEMORNA-CDS model for the generation of CDS based on the Transformer architecture (15).

### GEMORNA-CDS model demonstrates better codon usage regarding codon pairs, unwanted motifs, and in-cell stability

It has been shown that codon pairs significantly influence protein expression levels (16). Although two individual codons may each have moderate frequencies, their joint occurrence could be infrequent due to inhibitory interactions (17). Notably, natural sequences tend to have low frequencies of unwanted codon pairs, while sequences designed with widely used CAI-optimizing algorithms contain more unwanted codon pairs (**Fig. 2C**). In contrast, the AI-generated coding sequences (CDSs) exhibited a marked reduction in the incidence of unwanted codon pairs. Furthermore, we observe that AI-designed CDSs contain fewer slippery sites (18–19), reducing the +1 ribosomal frameshifting frequency (**Fig. 2D**). These findings suggest that the GEMORNA model has the capability to learn and apply codon usage patterns within a contextual framework.

The in-cell stability of mRNA sequences is also an important factor to consider when designing mRNA therapeutics and vaccines. Wu *et al*. investigated codon-dependent mRNA decay using an ORFome assay and used the previously described codon stability coefficient (CSC) as a measure of a codon’s contribution to mRNA stability (20). We found that AI-designed sequences significantly disfavor unstable codons, and lead to a higher correlation with CSC (**Fig. 2E**).

### GEMORNA-generated CDS demonstrated stronger and long-lasting expression

Next, we validated these results with cell-based assays, using a destabilized Firefly luciferase (denoted as Fluc2P) as the reporter. Three AI-generated coding sequences, labeled as GMR-FL1, GMR-FL2 and GMR-FL3, were compared against five other control CDSs: codon-optimized CDSs from commercial sources (GenScript and IDT), a CAI- and structure-jointly optimized CDS from a previous study (labeled as LinearDesign) (10), a deep-learning-based CDS from a biLSTM-CRF model (8), and a CDS from a commercially available vector (pGL4.11) (**Fig. 2F**). We observed that AI-designed CDSs exhibited a 4.7-fold improvement over biLSTM-CRF, and a 1.6-fold over the commercial pGL4.11 vector 24 hours after transfection (**Fig. 2G**). Given the rapid degradation of destabilized luciferase post-production, the Fluc activity ratio at 48 hours relative to 24 hours was used as a metric to evaluate the in-cell stability of mRNA constructs. AI-designed sequences exhibited higher Fluc activity ratio, suggesting they have greater in-cell stability (**Fig. 2H**). These experiments performed in in HepG2 cells showed consistent results with those generated in HEK293T cells (**Fig. SI 1**).

### GEMORNA-UTR: deep generative model for 5’ and 3’ UTR element design

In addition to mRNA CDSs, the UTR sequences that surround the CDSs play major role in the performance of mRNA therapeutics and vaccines by controlling expression levels and stabilities. Therefore, we developed GEMORNA-UTR model for the generation of UTRs.

### GEMORNA-UTR generated novel UTRs with better MRL and MFE

We first assessed the similarities between GEMORNA-generated UTRs and natural UTRs using a BLAST search. The Maximum Identity Score (MIS), defined as the longest aligned subsequence divided by the length of the query sequence, was employed as the measure of similarity. The MIS distributions for the 5’ UTRs show that most GEMORNA-generated UTRs exhibit low MIS values (**Fig. 3A**), suggesting that our AI-generate a significant number of novel UTRs that are dissimilar to natural UTRs. Furthermore, the ability to design 5’ UTRs with strong translation initiation would enable more potent mRNA products. Prior studies have shown that 5’ UTRs with higher MRL and MFE are more efficient in translation initiation (4, 13). The MRL and MFE distributions of our AI-generated 5’ UTRs shifted rightwards to higher values compared to the nature 5’ UTRs (**Fig. 3B**), which potentially leads to high efficiency translation initiation.

**Fig. 3:**
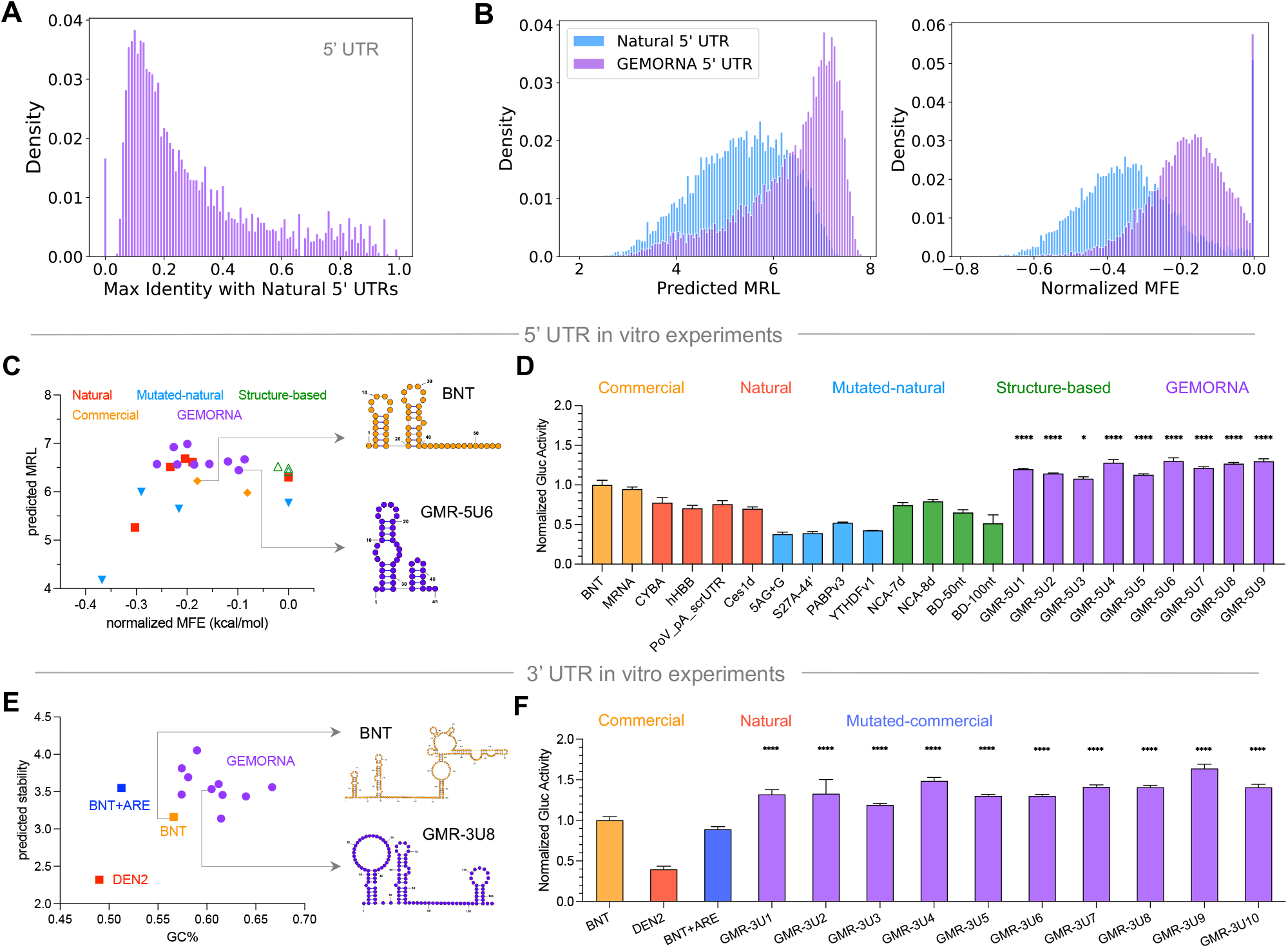
The *in silico* analysis and *in vitro* experimental results of UTR sequences generated by GEMORNA-UTR. **A)** Distributions of max identity between natural and GEMORNA 5’ UTRs; a small max identity indicates a low sequence similarity between our AI-generated UTRs and all natural UTRs in the reference database. **B)** MRL and MFE distributions of natural and GEMORNA-generated 5’ UTRs. **C)** Length-normalized MFEs and predicted MRLs for 5’ UTRs; the secondary structure of two representative 5’ UTRs are shown, one from BNT (commercial sequence) and the other designed by GEMORNA (GMR-5U6). **D)** Relative Gluc activities using GEMORNA 5’ UTRs compared to those using commercial, natural, and conventionally designed 5’ UTR counterparts. **E)** The GC percentage and predicted durability of 3’ UTRs; the secondary structure of two representative 3’ UTRs are shown, one from BNT and the other designed by GEMORNA (GMR-3U8). **F)** Relative Gluc activities of GEMORNA 3’ UTRs compared to control 3’ UTRs at 24 h. n.s.: no significance. \**P <* 0.05, \*\**P <* 0.01, \*\*\*\**P <* 0.0001.

### GEMORNA-generated UTRs demonstrated enhanced protein expression

We assessed 9 GEMORNA-designed UTRs for their ability to enhance protein expression compared to 14 high-performance 5’ UTRs as baselines, consisting of UTRs used in commercial mRNA drugs, natural, and optimized-natural UTRs (11; 12; 21) or structure-based rational design (12). A two-dimensional visualization of length-normalized MFE against mean ribosome loads (MRL) for the UTRs is shown in **Fig. 3C**. MRLs were calculated by our deep learning-based predictive model PRED-5UTR, which exhibits high correlation between experimental and predicted MRLs (**Fig. SI 2**). Normalized MFE is the MFE of a given 5’ UTR divided by its length, with a higher value indicating less secondary structure. Our AI-generated 5’ UTRs had high predicted MRLs, and relatively high normalized MFEs. **Fig. 3D** shows the Gluc activity at 48 h in HEK293T. Among the 14 baselines, the 5’ UTRs used in BNT162b2 and mRNA-1273 COVID mRNA vaccines, denoted as BNT and MRNA, showed the highest Gluc activities. Nevertheless, we observed that GEMORNA 5’ UTRs exhibited similar or higher Gluc activities than BNT and MRNA.

We subsequently evaluated the performance of GEMORNA in 3’ UTR design. We chose 10 AI-derived 3’ UTRs against another three baselines, including BNT162b2 3’ UTR (denoted as BNT in **Fig. 3E** and **F**), a BNT162b2 3’ UTR variant built by adding an AU-rich element (denoted as BNT+ARE), and a natural one from dengue virus (denoted as DEN2) that increases viral protein expression (11). Similar to the 5’ UTR analysis above, we first visualized these thirteen 3’ UTRs in a two-dimensional plot, with the *x*-axis being the GC percentage and *y*-axis being the predicted in-cell stability (**Fig. 3E**). The stabilities were calculated by our predictive model, PRED-3UTR, which was trained and tested on experimental data. Our AI-designed 3’ UTRs were in the zone of higher GC percentage and predicted stability. GEMORNA 3’ UTRs exhibited higher Gluc activity in both HEK293T and Jurkat cells at 48 hours (**Figs. 3F and SI 3**) and 24 hours (**Fig. SI 4**) post transfection. These results showed an interesting correlation between gene expression levels and the predicted stability of 3’ UTRs (22), suggesting a potential approach for 3’ UTR optimization.

## Full-length mRNA design with GEMORNA-generated CDSs and UTRs

In previous sections, we showed that both GEMORNA-generated CDSs and UTRs lead to higher *in vitro* translation capacity compared to a variety of baseline sequences. Next, we investigated whether full-length mRNA designs derived from GEMORNA elements could further enhance translation capacity, and lead to better immunogenicity in mRNA vaccines.

### GEMORNA full-length mRNAs achieved higher protein expression and durable expression

We first designed full-length mRNAs to express the Fluc2P reporter. We designed five mRNA constructs with GEMORNA-generated CDSs and UTRs, and compared these to two benchmark mRNAs: the first consisting of an IDT-designed CDS with natural alpha globin (AG) UTRs (labeled as Benchmark-FL1), and the second consisting of the pGL4.11 CDS with BNT UTRs (labeled as Benchmark-FL2). We observed that all of our AI-designed full-length sequences achieved significantly higher Fluc activity than the two benchmarks, with a more pronounced enhancement at 48 h compared to 24 h (**Fig. 4A**). The higher gene expression ratio of 48 h to 24 h indicated our AI-designed mRNAs might have longer half-lives compared with the benchmark mRNAs. Remarkably, the most effective full-length design, GMR-FL-F5, achieved an 8.2-fold increase in Fluc activity at 48 h when compared to the more robust Benchmark-FL2. The protein expression improvements were also observed in HepG2 cells (**Fig. 4B**).

**Fig. 4:**
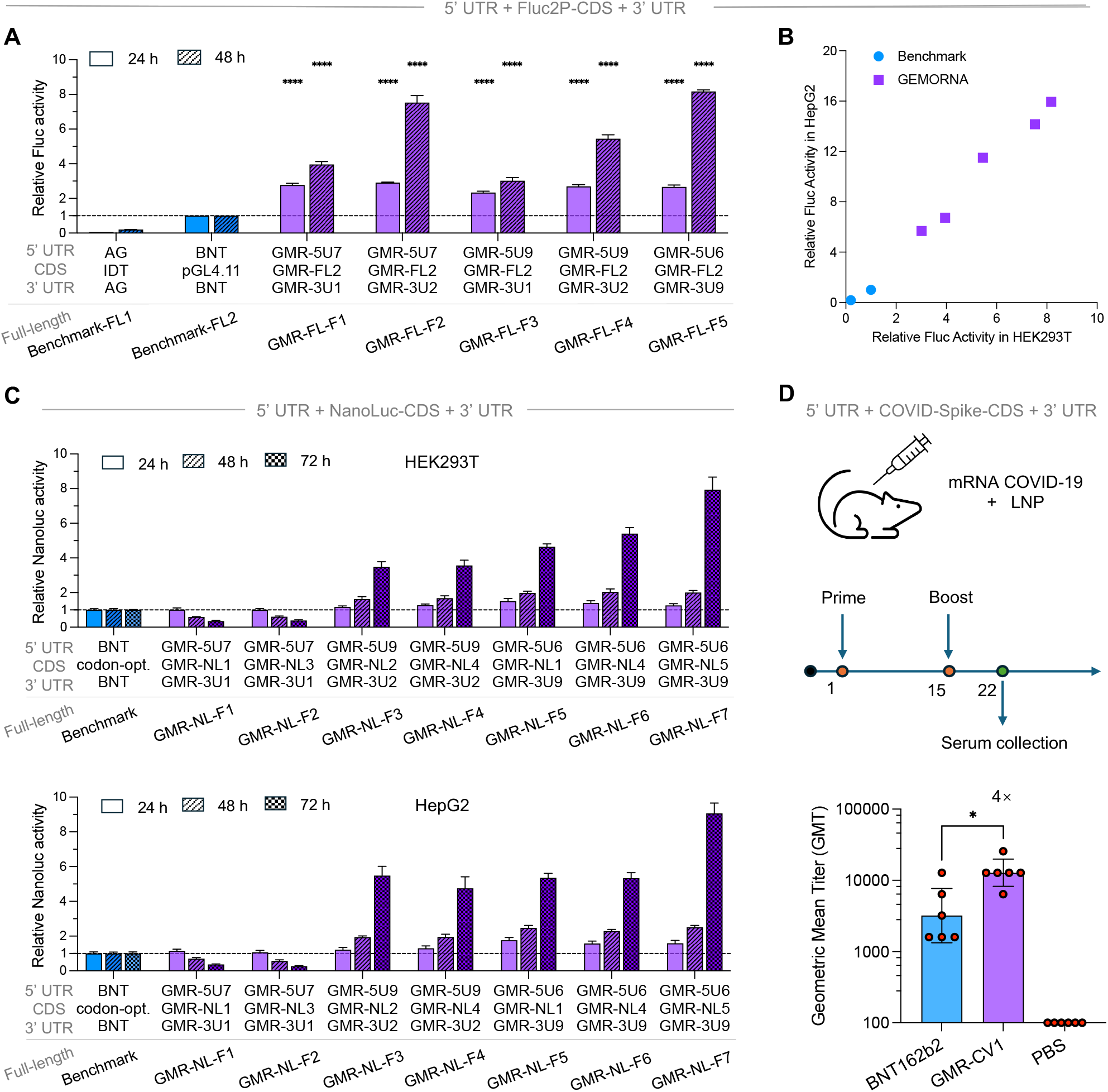
By combining GEMORNA-generated CDS and UTRs, we designed full-length mRNAs with stronger *in vitro* protein expression and *in vivo* immunogenicity compared with conventionally designed counterparts and commercial products. **A)** Relative Fluc activities of seven full-length mRNAs encoding the Fluc2P reporter with different UTRs and CDSs. **B)** Relative Fluc activities at 48 h in HEK293T and HepG2 cells. **C)** Relative NanoLuc activities at 24, 48 and 72 hours in HEK293T and HepG2 cells. **D)** Head-to-head comparisons of anti-spike IgG triggered in mice between our full-length COVID-19 mRNA vaccine sequence (GMR- CV1) and the commercial BNT162b2 counterpart. Bar plots are presented with mean *±* s.d. for *in vitro* data and geometric mean*±* geometric s.d. for *in vivo* data. \**P <* 0.05, \*\*\*\**P <* 0.0001.

We subsequently validated GEMORNA platform with a second commonly used reporter protein, NanoLuc. Seven AI-enabled mRNAs were compared to a benchmark sequence which is consisting of codon-optimized CDS and BNT UTRs. Five out of seven AI designs exhibited higher NanoLuc expression levels, with observed improvements increasing from 24 h to 72 h (**Fig. 4C**). The highest fold-changes over benchmark sequence were 7.9× and 9.1× in HEK293T and HepG2 cells, respectively.

### GEMORNA-designed mRNA vaccine demonstrated higher immunogenicity

Next, we used GEMORNA to generate a therapeutic mRNA coding the COVID-19 spike protein, and tested its immunogenicity in mice against BNT162b2. Lipid nanoparticle (LNP) encapsulated mRNAs were injected into six BALB/c mice for each sequence. We injected two doses, spaced apart by two weeks, and collected the serum of immunized mice one week after the last dose to measure SARS-CoV-2-specific IgG antibody levels. (**Fig. 4D**). These results showed that our GEMORNA-derived full-length mRNA (labeled as GMR-CV1) induced a 4-fold higher antibody titer than BNT162b2 at the same LNP-mRNA dose level.

### Circular RNA design with GEMORNA

Attributed to their circular topology, circular RNAs (circRNAs) exhibit increased resistance to exonucleases compared to their linear RNA counterparts (23). This property contributes to their enhanced in-cell stability relative to linear RNAs; however, efforts to further strengthen this advantage are still of great interest (21). Conversely, circRNAs necessitate a cap-independent mechanism for translation due to the absence of a 5’-cap, a process that is generally less efficient than the cap-dependent translation observed in linear mRNAs (24). Therefore, it is highly intriguing to investigate whether GEMORNA is extensible to circRNA design, generating circRNAs with further-enhanced durability, improved expression levels, and strong efficacy.

### GEMORNA generated circRNAs demonstrated high and long-lasting expression

Initial tests using Fluc2P assays revealed that GEMORNA-generated CDSs outperformed conventionally designed CDSs within the circular system (**Fig. SI 5**). In addition to CDS optimization, a previous study has demonstrated enhanced circRNA expression with optimized topology, screened UTRs and engineered IRES (21). We hypothesized that the performance of circRNA could be further elevated by our AI-generated CDSs and mined IRESs, even with a simplified topology (**Fig. 5A and B**). To verify this, we conducted additional experiments using NanoLuc assays. The best-known benchmark (labeled as Chen et al.) was a systematically optimized circRNA, consisting of PABP 5’ UTR, engineered HRV-B3 IRES, codon-optimized CDS and hHBA1 3’ UTR (21); our designs employed different AI-generated CDSs combined with IRESs mined from viral UTRs. Compared to the benchmark, all our designs exhibited significantly higher accumulated NanoLuc expression, with the maximum increase reaching 4.7-fold over the benchmark in HEK293T cells (**Fig. 5C**). More importantly, GEMORNA sequences exhibited strong expression durability. Our design, GMR-NL3, maintained a high NanoLuc activity over 144 h, achieving a 3-fold increase in NanoLuc activity at 72 h compared to 24 h. In contrast, the NanoLuc activity of the benchmark continuously declined after 48 h. We observed similar results in HepG2 cells (**Fig. SI 6**).

**Fig. 5:**
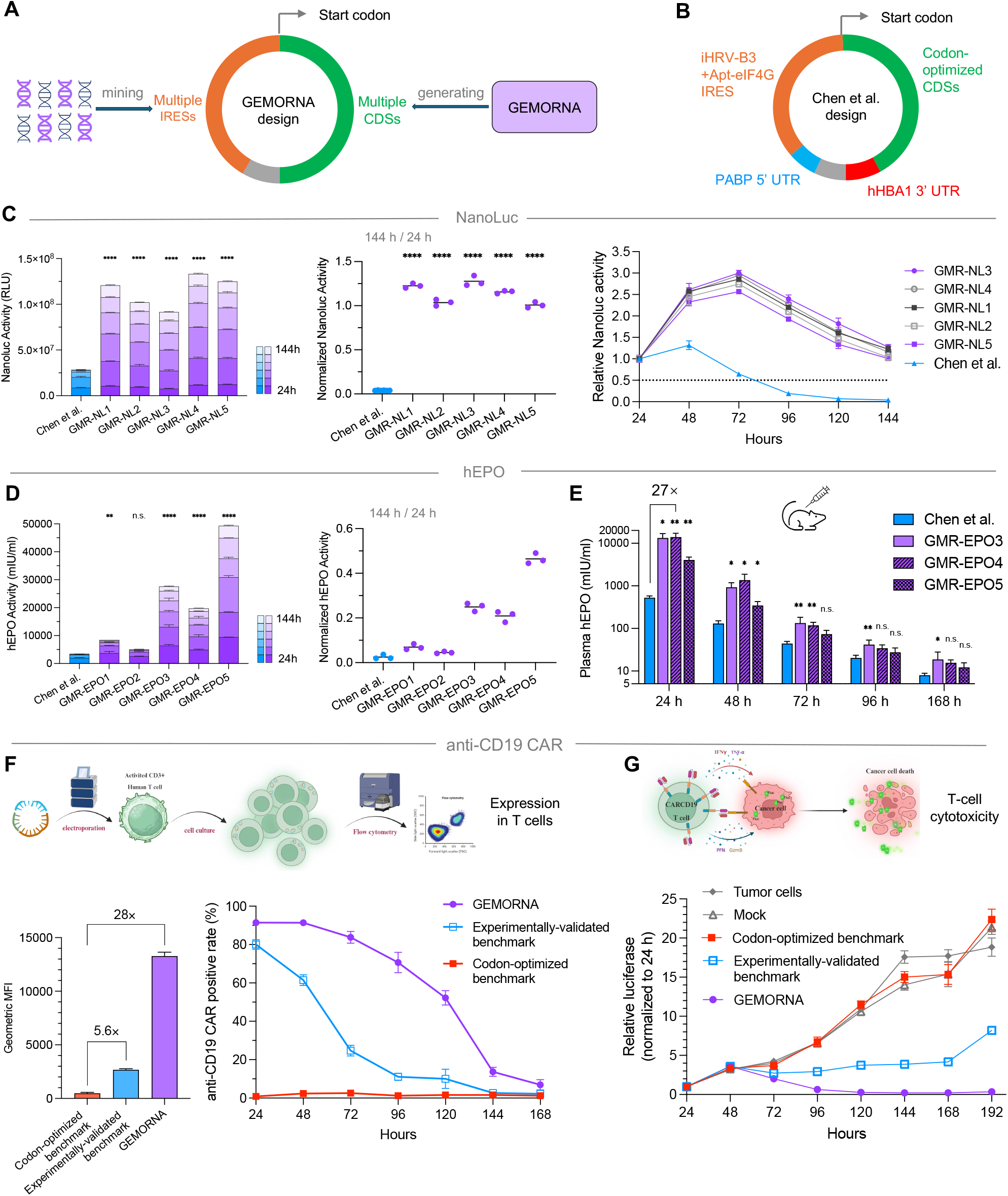
The GEMORNA platform is able to generate circRNAs with improved expression level, durability and druggability compared with conventionally designed and patented counterparts. **A)** Our AI-enabled circRNA topology integrates with mined IRESs and AI-generated CDSs. **B)** The optimized circRNA topology proposed by Chen et al. (21) which uses screened or optimized UTRs, IRES and CDSs; this topology was used as a benchmark compared with our GEMORNA-enabled design approach. **C)** Comparison of NanoLuc expression levels and expression durability between GEMORNA circRNAs and a benchmark circRNA in HEK293T cells. Accumulated NanoLuc within 144 h, ratios of NanoLuc product at 144 h to product at 24 h, and relative NanoLuc activities from 24 h to 144 h are shown. **D)** Comparison between GEMORNA circRNAs and a benchmark circRNA expressing hEPO protein in HEK293T cells; accumulated hEPO protein levels over 144 h are shown, and expression level ratios of 144 h to 24 h are also shown. **E)** hEPO protein expression in mice. **F)** Expression levels and durability of three circRNAs expressing anti-CD19 CAR in T cells. **G)** Comparisons of *in vitro* T-cell killing of NALM-6 cells (E:T=1:3) between GEMORNA circRNAs and two control mRNAs. “Tumor cells”: no T-cell treatment. “Mock”: T-cell treatment without anti-CD19 CAR circRNAs transfected. All these circRNAs encode anti-CD19 CAR. n.s.: no significance. \**P <* 0.05, \*\**P <* 0.01, \*\*\*\**P <* 0.0001.

We further evaluated circRNAs encoding human erythropoietin (hEPO) to determine whether the observed improvements extend to therapeutic proteins. Similarly, the benchmark sequence was constructed with elements showed in **Fig. 5B**, and its CDS was a codon-optimized one from previous studies (21). **Fig. 5D** illustrates the hEPO expression from 24 h to 144 h in HEK293T cells, showing that our best design exhibited 13.8-fold increase in accumulated expression compared to benchmark. Additionally, GEMORNA sequences showed a superior ratio of expression levels at 144 h compared to 24 h, with the highest ratio being 46.5%, in contrast to the benchmark’s 2.5%. Similar results were observed in HepG2 cell lines (**Fig. SI 7**).

Subsequently, we selected three GEMORNA circRNAs, GMR-EPO3, GMR-EPO4 and GMR-EPO5, which exhibited strong translation capacity *in vitro*, for an *in vivo* evaluation. LNP-circRNAs were injected into mice, and serum samples were collected between 24-168 hours post-injection to measure expression levels. All three GEMORNA circRNAs demonstrated significantly higher *in vivo* hEPO expression levels at 24 h, with the most effective circRNA showing a 27-fold increase compared to the benchmark (**Fig. 5E**). Additionally, our AI- designed sequences maintained relatively higher expression levels from 24 h to 168 h, confirming their long- lasting expression property *in vivo*.

### GEMORNA circRNAs demonstrated enhanced efficacy for non-viral CD19-CAR T therapy

While CAR-T cell therapy has achieved great success in treating hematologic malignancies, the manufacturing process remains time-consuming and costly. Given the success of mRNA vaccines, there is increasing interest in extending the mRNA therapeutic platform to *in vivo* CD19-CAR T therapy. CircRNAs, which offer extended protein expression durability, were used in producing therapeutic CAR-T cells expressing CD-19 CAR within the human body. Here we conducted non-viral CD19-CAR T experiments *in vitro* to assess the durability and the therapeutic potential of our AI-enabled circRNAs for CAR-T therapy.

We first constructed two baseline constructs, one denoted as “Codon-optimized benchmark”, with the coding region designed using a conventional codon optimization approach, and the other denoted as “Experimentally validated benchmark”, a patented circRNA sequence showed strong gene expression and tumor killing efficacy (25). The baselines and GEMORNA-designed circRNA were then delivered into primary human T cells. Our circRNA exhibited significant increases in gene expression, surpassing “Codon-optimized benchmark” by 28- fold at 24 hours post-electroporation. In contrast, “Experimentally validated benchmark” showed a 5.6-fold increase relative to “Codon-optimized benchmark” (**Fig. 5F**). Additionally, 50% of the cells with GEMORNA-designed circRNA were still CD-19 CAR-positive 120 hours post-electroporation, compared to less than 72 hours for the Benchmark circRNAs (**Fig. 5F**). We also confirmed our AI-enabled circRNA showed significant improvements in gene expression level and tumor cell cytotoxicity over a linear mRNA control (**Fig. SI 8**). We further investigated the cytotoxic efficacy of circRNA candidates in primary human T cells against NALM- 6, human cancerous B cells expressing CD19 (26), were incubated with NALM-6 target cell at effector:target (E:T) ratio of 1:6, and co-cultivated for 7 days at 37°C. As shown in **Figs. 5G and SI 9**, our high-expression and long-lived circRNAs significantly inhibited the growth of NALM-6 cells, while the control sequence showed virtually no cytotoxic effect. The benchmark sequence “Experimentally validated benchmark” was also less efficient than our design.

In summary, our results indicate that the high and durable gene expression facilitated by GEMORNA effectively enhanced CD19-CAR T cell cytotoxicity *in vitro*. We inferred that our platform could improve the therapeutic potential of circRNAs for *in vivo* CAR-T therapy.

## Discussion

mRNA therapeutics have been recognized as a promising next-generation modality. Nevertheless, low protein expression levels and short duration of protein expression pose significant challenges to their broader application in fields such as immuno-oncology and gene replacement therapy. To address these limitations, we proposed GEMORNA, generative models designed to enhance mRNA translation capacity, and experimentally confirmed that GEMORNA-derived elements contribute to the development of superior mRNA drug molecules. Compared to previous mRNA optimization methods, our strategy confers distinct advantages. First, GEMORNA-generated elements exhibit enhanced translation capacity with m1Ψ modification, which is pivotal to the success of marketed mRNA drugs (18). Second, our experiments confirmed that GEMORNA is extensible to the design of circular RNAs, resulting in substantial improvements in long-lasting expression.

Third, unlike prior studies that trained RNA language models primarily for predictive tasks (27; 28), GEMORNA was conceived as deep generative models for directly generating high-quality mRNA elements. To our best knowledge, GEMORNA represents the first generative model for the design of full-length mRNAs. Finally, GEMORNA-derived sequences were benchmarked against commercial products and the most advanced designs in the literature, exhibiting significant improvements in expression levels, durability and immunogenicity. In conclusion, our generative AI-based model constitutes a novel and efficacious strategy for mRNA sequence design, with the potential to notably advance the development of mRNA vaccines and therapeutics.

## Acknowledgments

The authors thank other current or former employees, contractors, or consultants of Raina Biosciences who provide advice or assistance on the studies described in the manuscript.

## Materials and Methods

### Plasmid construction and RNA synthesis

The plasmids used in this study were built using restriction enzyme cloning and Gibson assembly. The in vitro transcription (IVT) templates were PCR-amplified. Subsequently, the mRNA and circular RNA (circRNA) were synthesized as described previously (10; 23).

### Cell culture and transfections

HEK293T and HepG2 cells were cultured in DMEM (Thermo Fisher) supplemented with 10% FBS and 100 units/mL penicillin and streptomycin. Jurkat cells and NALM-6 expressing firefly luciferase- GFP (NALM-6) were cultured in RPMI 1640 (Thermo Fisher) supplemented with 10% FBS and 100 units/mL penicillin and streptomycin. For HEK 293T and HepG2 cells, the cells were plated on 96-well plates one day before transfection, and the transfection was performed with Lipofectamine MessengerMax (Thermo Fisher) according to the manufacturer’s manual. For Jurkat and NALM-6 cells, the cells were seeded at the time of transfection with Lipofectamine MessengerMax according to the manufacturers’ manuals.

### Luciferase assay

Firefly luciferase activity was measured with the kit (Promega) according to the manufacturer’s instructions. Gaussia Luciferase activity was measured with the kit (Beyotime) according to the manufacturer’s instructions. NanoLuc luciferase activity was with the kit (Promega) measured according to the manufacturer’s instructions.

### Flow cytometry

Expanded T cells were stained for CD19-CAR expression with biotinylated human CD19 and PE Streptavidin combination at indicated time points. Cells were also co-stained with anti-human CD3, anti-human CD4 and anti-human CD8 (Biolegend), then assayed using a BD LSRFortessa flow cytometer (BD Biosciences).

### hEPO ELISA assay

Erythropoietin was detected by ELISA (Elabscience) essentially according to the manufacturer’s instructions except cell culture supernatant at indicated time points post transfection were used, and samples were diluted 1:200 before use.

### LNP encapsulation of the RNAs

The mRNAs were encapsulated with lipid nanoparticles (LNPs). First, the RNAs were diluted with PNI Formulation Buffer (Precision NanoSystems, #NWW0043) to a final concentration of 170 µg/ml. Then, the lipids were mixed with the mRNA solution at the volume ratio of 1:3 through the Ignite NxGen Cartridge (Precision NanoSystems, #NIT0002) using NanoAssemblr Ignite (Precision NanoSystems). Then the LNP-mRNA formulations were diluted by 40-fold with 1*×*PBS buffer (pH 7.2-7.4) and concentrated via ultrafiltration with Amicon Ultra Centrifugal Filter Unit (Millipore). The concentration and encapsulation efficiency of mRNAs were measured by the Quant-it RiboGreen RNA Assay Kit (Invitrogen, #R11490). The size of LNP-mRNA particles was measured using dynamic light scattering on a Malvern Zetasizer Nano-ZS 300 (Malvern).

### *In vivo* delivery of mRNA and circRNA

For mouse vaccination, groups of 6- to 8-week-old female BALB/c mice were intramuscularly immunized with LNP-mRNAs as indicated or a placebo (LNP only) using a 1-ml sterile syringe, and 2 or 3 weeks later, a second dose was administered to boost the immune responses. The sera of immunized mice were collected for the detection the SARS-CoV-2-specific IgG antibody by ELISA assay as described below. All experiments using mice were conducted under the ethical regulations and were approved by local ethical committees.

For *in vivo* EPO expression studies, groups of 6- to 8-week-old female BALB/c mice were injected with 15 micrograms of LNP-circRNAs once on day 0, intravenously via the tail vein as indicated or a placebo (LNP only) using a 1-ml sterile syringe. Sera of LNP-circRNA injected mice were collected at different time points and the EPO concentrations were measured by ELISA assay. All experiments using mice were conducted under the ethical regulations and were approved by local ethical committees.

### IgG antibody ELISA

All immunized mouse serum samples were heat-inactivated at 56 °C for 30 min before use. The SARS-CoV-2-specific IgG antibody was measured a commercial ELISA kit (Vazyme) according to the manufacturer’s instructions. Briefly, serial 3-fold dilutions of heat-inactivated sera, starting at 1:100, were added to 96-well plates (100 µl/well) coated with recombinant SARS-CoV-2 RBD antigen and incubated for 60 min at 37 °C. After three washes with the wash buffer, horseradish peroxidase HRP-conjugated rabbit anti-mouse IgG was added to the plates and incubated at 37 °C for 30 min. Then, the plates were washed three times with the wash buffer. After washing, the HRP substrates and stop solution were added, and the absorbance at 450 nm was measured with 630 nm as a reference with a Synergy H1 Microplate Reader (BioTek).

### Generation of CD19-CAR T-cells by electroporation

*In vitro* expanded human primary T cells were subjected to electroporation per the manufacturer’s instructions using *in vitro* transcribed mRNAs or circRNAs. Prior to electroporation with a Gene Pulser Xcell Eukaryotic System (Bio-Rad), expanded T cells were counted, assessed for viability with Trypan Blue, and washed twice with Dulbecco’s Phosphate-Buffered Saline (Solarbio). Cells were then resuspended at a density of 1 *×* 10^7^ cells/ml in Gene Pulser Electroporation Buffer (Bio-Rad). Aliquots of the cell/electroporation buffer mix were transferred into 1.5 ml microcentrifuge tubes (100 µl per condition). *In vitro* transcribed circRNAs was added to each mix. The mixture was homogenized by pipetting up and down, then transferred into a sterile Gene Pulser/MicroPulser Electroporation Cuvette with 0.2 cm gap (Bio-Rad) and pulsed with voltage 220 V and pulse lengths of 2 ms using the square waveform pulse. After pulsing, cells were immediately transferred into a 24-well plate containing 900 µl prewarmed T Cell Expansion Medium.

### *In vitro* cytotoxicity of CD19-CAR T-cells

Luciferase-expressing NALM-6 cells were plated into 96-well plates and co-cultured with the indicated ratio of CD19-CAR T cells for 7 days. Cells were washed with DPBS before being lysed and assayed for luciferase luminescence according to the manufacturer’s recommendations (Vazyme, DD1204-02) on a BioTek plate reader. A decrease in luciferase indicates the FAP expressing HEK293T target cells were eliminated by functional CD19-CAR T cells. Killing efficacy was represented as Relative Luminescent Units (RLU).

### Statistical analysis

All quantitative data are presented as mean ± standard deviation (s.d.). Statistical significance test was assessed using a one-way ANOVA followed by Dunnett’s multiple comparisons test, compared to the labeled control.

---

**Fig. SI 1.**
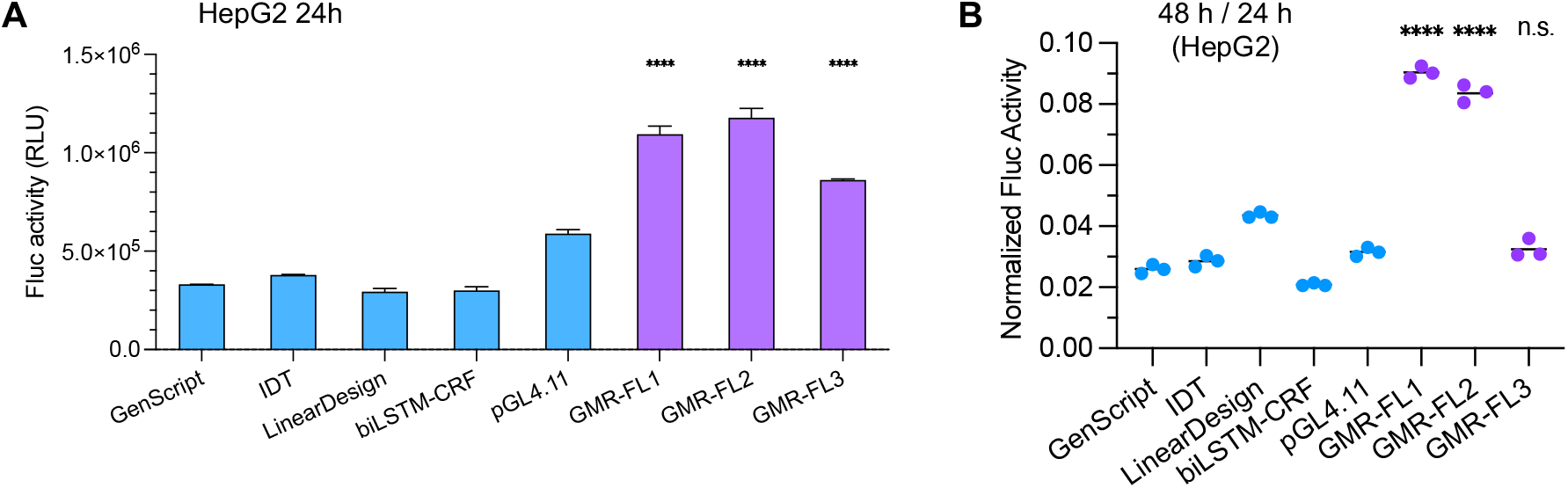
Comparisons of expression levels and duration between GEMORNA CDSs and control CDSs in HepG2 cells. **A)** Fluc activity at 24 h. **B)** Ratio of expression level at 48 h to expression level at 24 h. n.s.: no significance. \*\*\*\**P* < 0.0001.

**Fig. SI 2.**
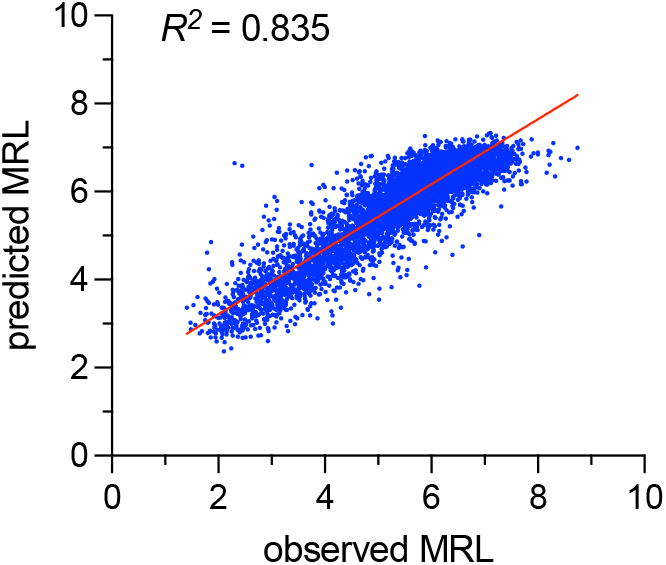
Correlation between observed mean ribosome load (MRL) and PRED-5UTR model predicted MRL.

**Fig. SI 3.**
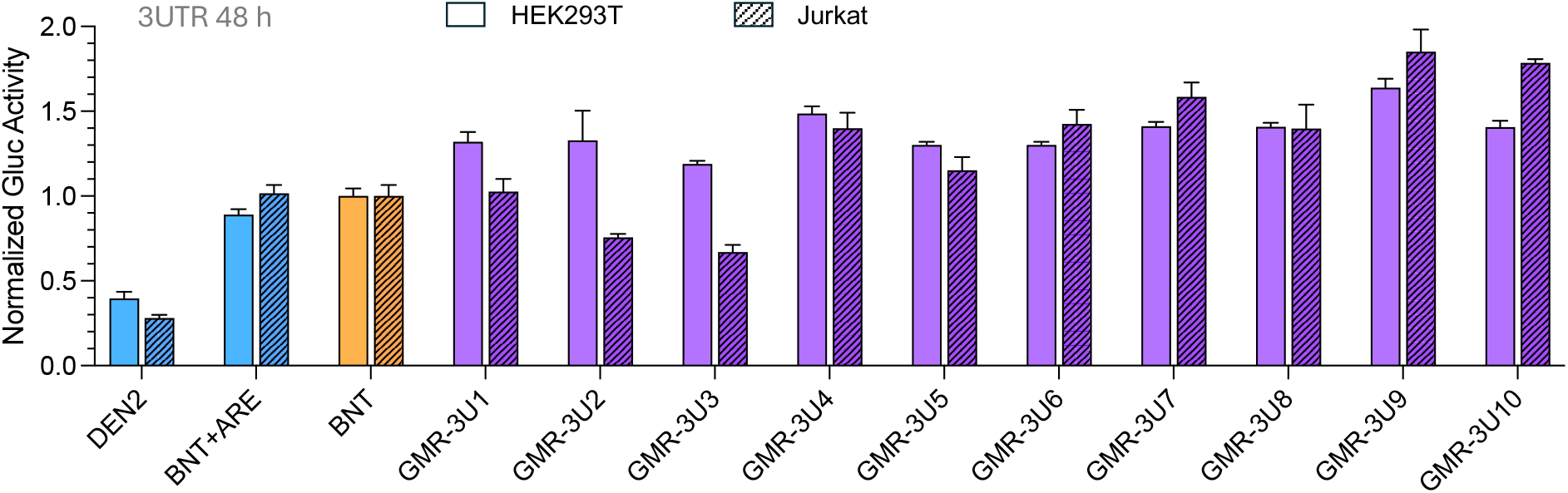
Relative Gluc activity of GEMORNA 3’ UTRs compared to baselines at 48 h in Jurkat cells.

**Fig. SI 4.**
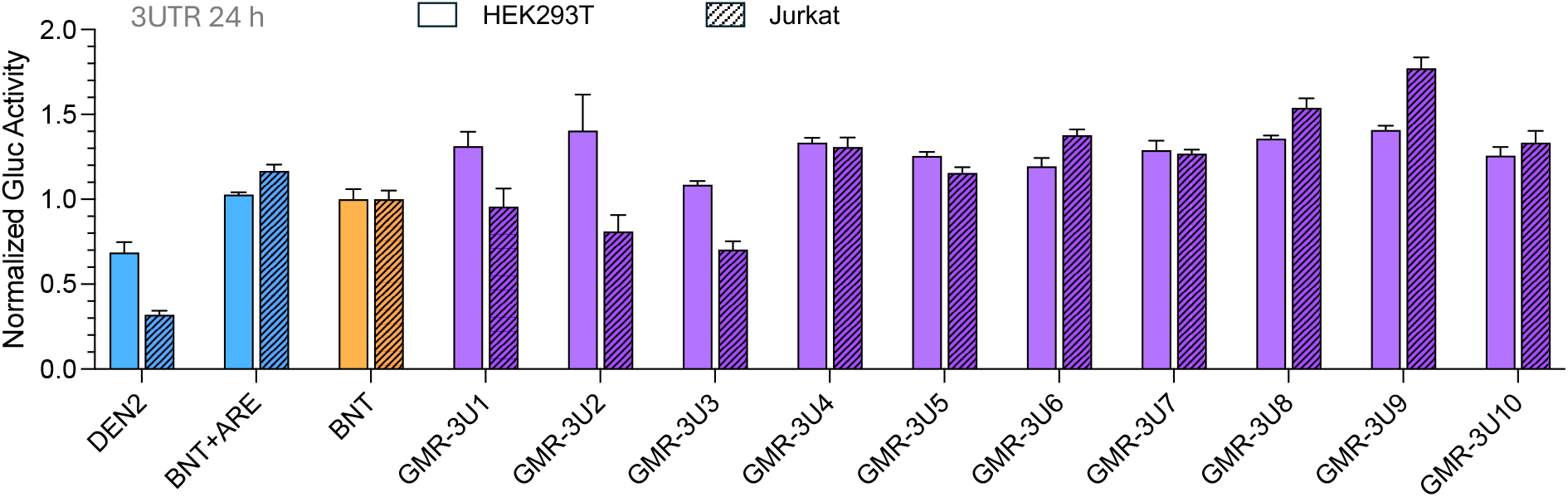
Relative Gluc activity of GEMORNA 3’ UTRs compared to baselines at 24 h. Data for both HEK293T and Jurkat cell lines are shown.

**Fig. SI 5.**
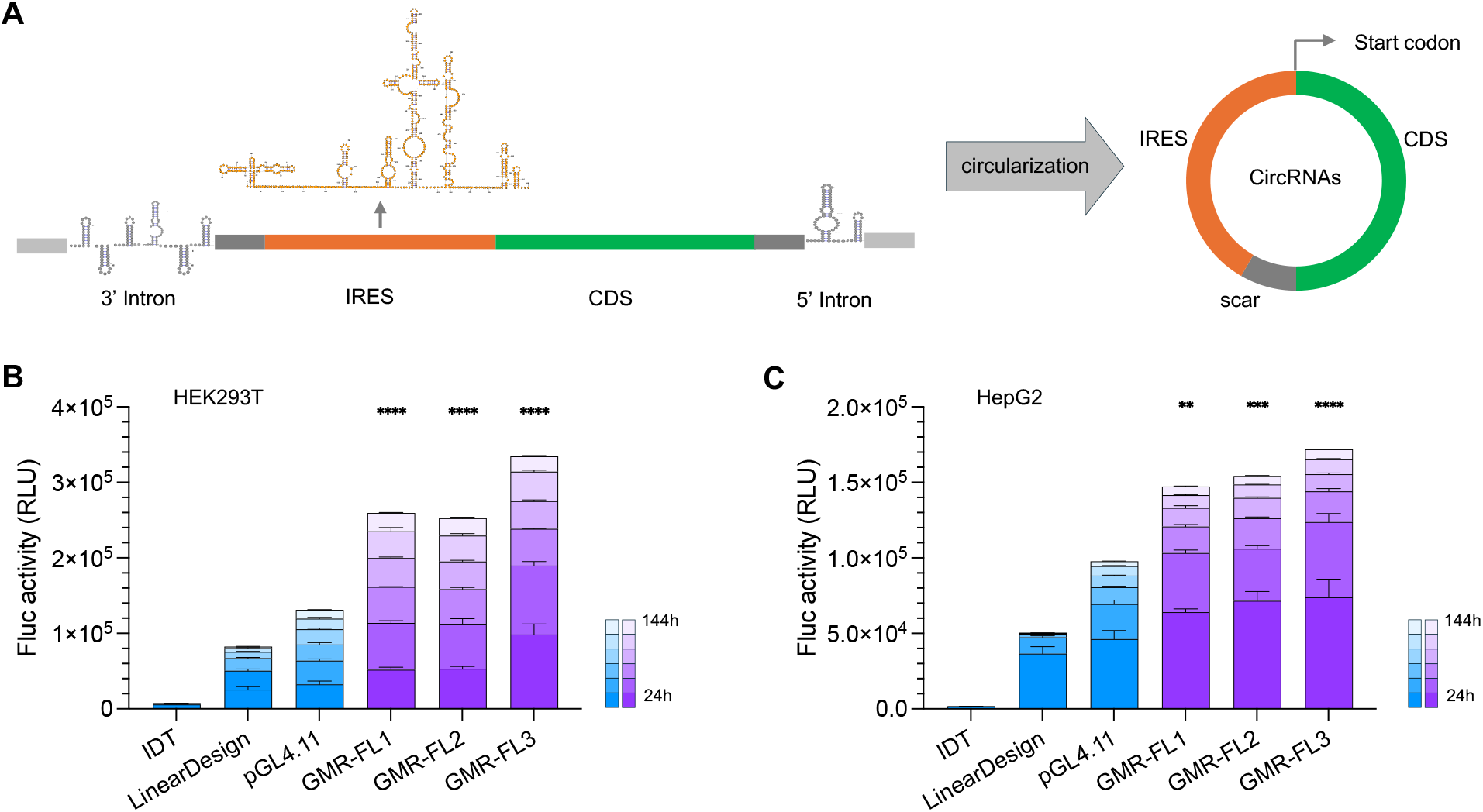
The GEMORNA platform is able to generate circRNA CDSs with enhanced expression levels and durability. **A)** Visualization of circular RNA components and circularization process. For the circRNA constructs, we followed the intron-based circularization method (23) and utilized the initial ribosome entry site (IRES) from the coxsackievirus B3 (CVB3) genome. **B)–C)** Fluc2P expression levels in HEK293T (**B**) and HepG2 (**C**) cell lines. n.s.: no significance. \*\**P <* 0.01, \*\*\**P <* 0.001, \*\*\*\**P <* 0.0001.

**Fig. SI 6.**
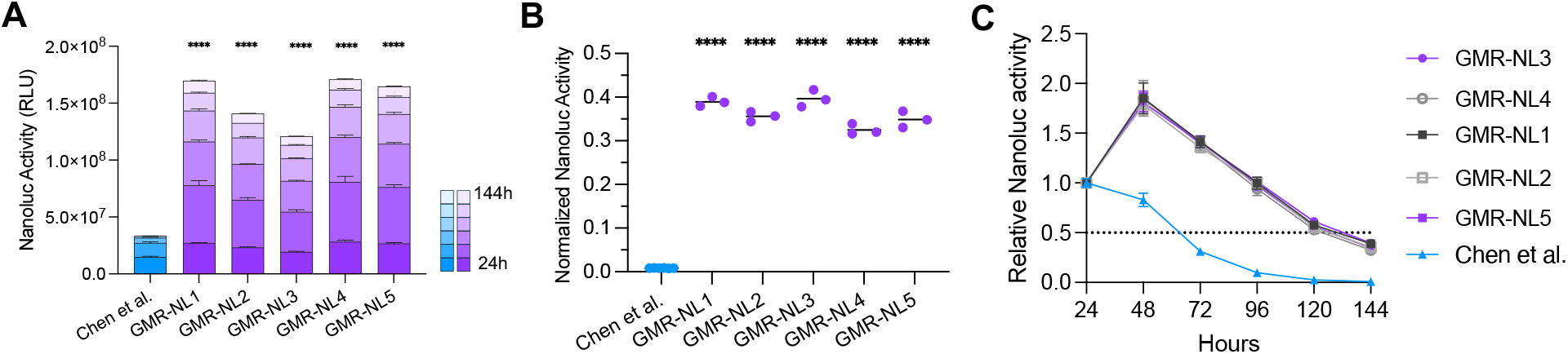
Expression levels and durability of NanoLuc circRNAs in HepG2 cells. **A)** Comparison of accumulated NanoLuc between GEMORNA circRNAs and the benchmark circRNA. NanoLuc activities were measured every 24 hours. **B)** Ratios of NanoLuc product at 144 h to product at 24 h. **C)** Relative NanoLuc activities from 24 h to 144 h after transfection; the data were normalized to NanoLuc activity at 24 h. n.s.: no significance. \*\*\*\**P <* 0.0001.

**Fig. SI 7.**
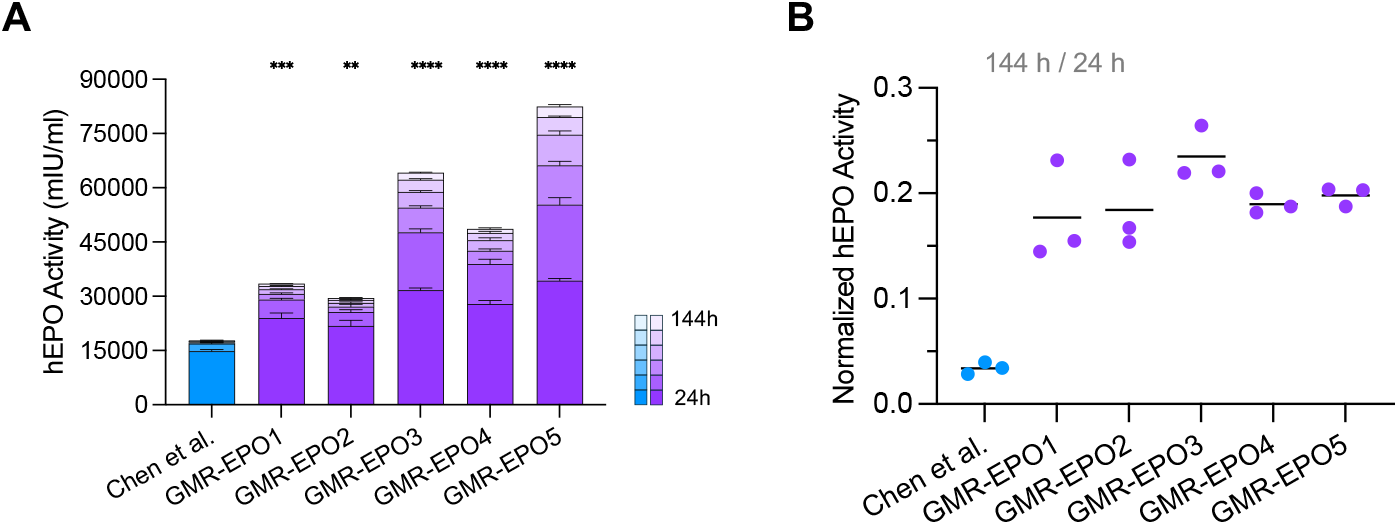
Expression levels and durability of hEPO circRNAs in HepG2 cells. **A)** Comparison of accumulated hEPO between GEMORNA circRNAs and Chen et al. benchmark. **B**) hEPO expression level ratios of 144 h to 24 h. \*\**P <* 0.01, \*\*\**P <* 0.001, \*\*\*\**P <* 0.0001.

**Fig. SI 8.**
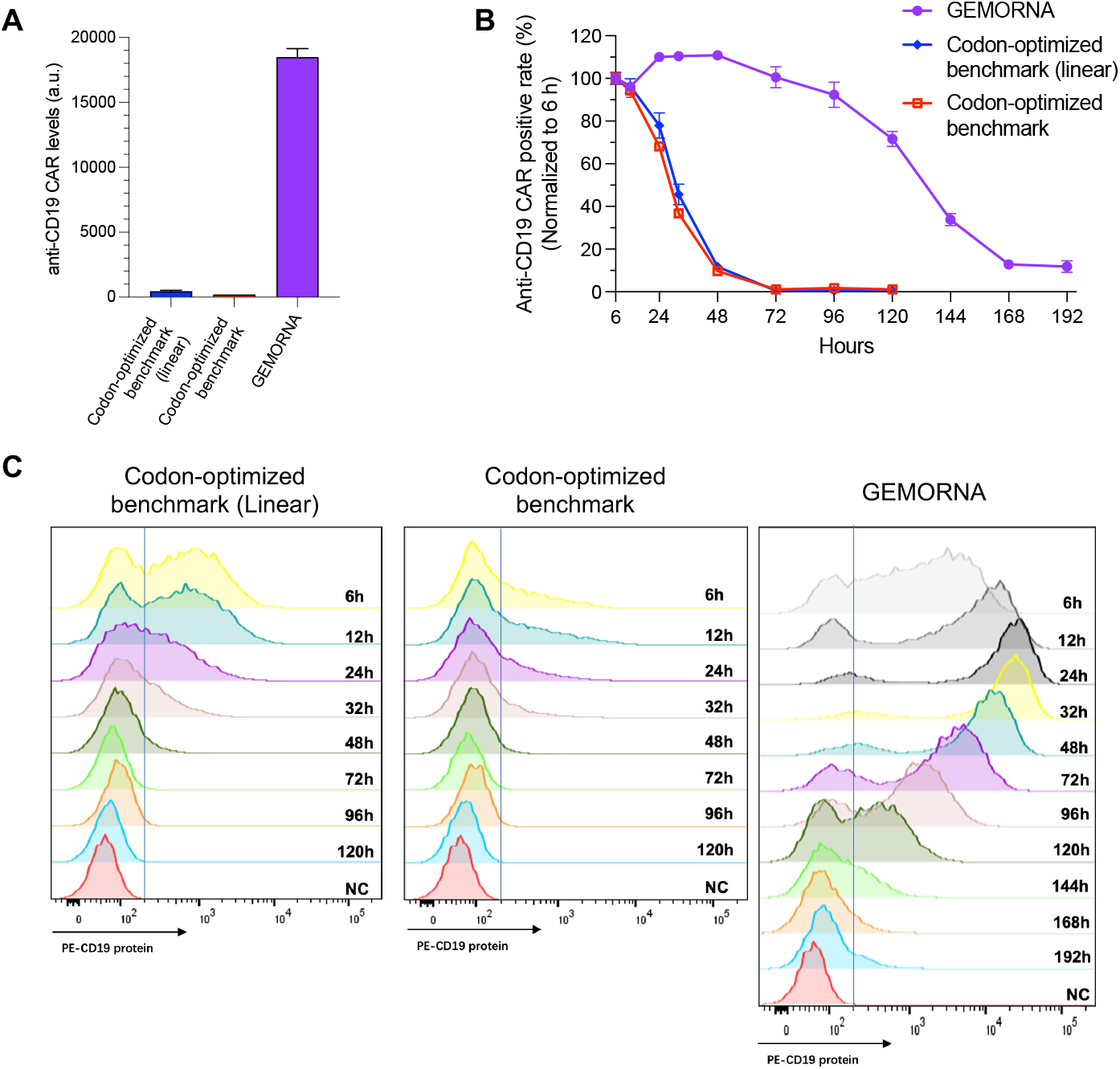
Comparisons of anti-CD19 CAR expression levels between GEMORNA circRNA and two conventionally designed benchmark RNAs. The two benchmarks have the same CDS, but one is linear RNA, denoted as “codon-optimized benchmark (linear)” and the other is circRNA, denoted as “codon-optimized benchmark”. **A)** Expression levels of the three sequences at 24 h after electro-transfected into Jurkat cells. **B)** Changes in the anti-CD19 CAR positive rate over 192 h. Electroporation efficiency was normalized by positive rate at 6 hours post electroporation. **C)** Detailed anti- CD19 CAR expression from 6 h to 192 h after electro-transfected into Jurkat cells.

**Fig. SI 9.**
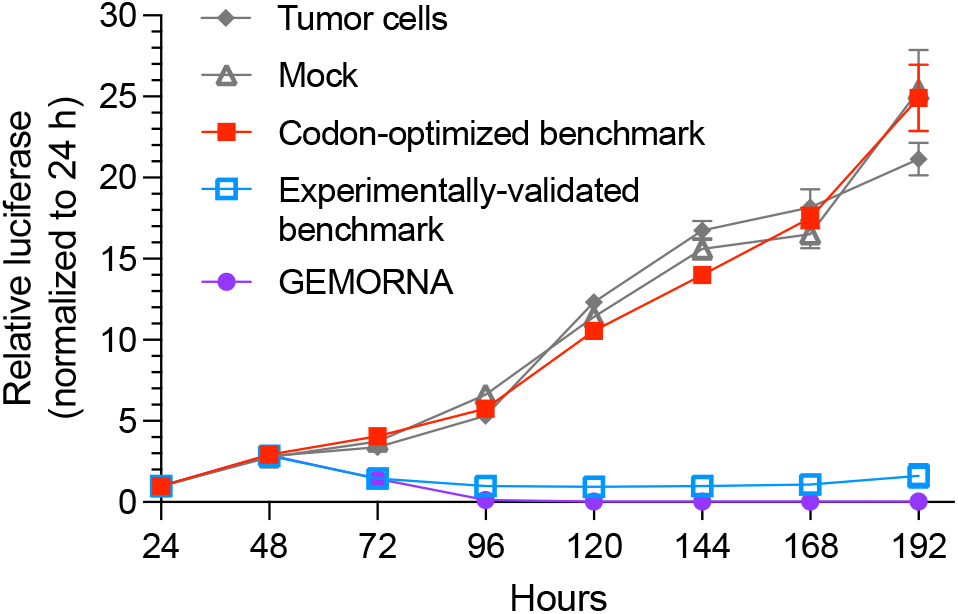
T-cell cytotoxicity experiment with a effector:target (E:T) ratio of E:T=1:3.

